# Gene amplification mutations originate prior to selective stress in *Acinetobacter baylyi*

**DOI:** 10.1101/2022.08.23.504990

**Authors:** Jennifer A. Herrmann, Agata Koprowska, Tesa J. Winters, Nancy Villanueva, Victoria D. Nikityuk, Feini Pek, Elizabeth M. Reis, Constancia Z. Dominguez, Daniel Davis, Eric McPherson, Staci R. Rocco, Cynthia Recendez, Shyla M. Difuntorum, Kelly Faeth, Mario D. Lopez, Habeeba M. Awwad, Rola A. Ghobashy, Lauren Cappiello, Ellen L. Neidle, Semarhy Quiñones-Soto, Andrew B. Reams

## Abstract

The controversial theory of adaptive amplification states gene amplification mutations are induced by selective environments where they are enriched due to the stress caused by growth restriction on unadapted cells. We tested this theory with three independent assays using an *Acinetobacter baylyi* model system that exclusively selects for *cat* gene amplification mutants. Our results demonstrate all *cat* gene amplification mutant colonies arise through a multistep process. While the late steps occur during selection exposure, these mutants derive from low-level amplification mutant cells that form before growth-inhibiting selection is imposed. During selection, these partial mutants undergo multiple secondary steps generating higher amplification over several days to multiple weeks to eventually form visible high-copy amplification colonies. Based on these findings, amplification in this *Acinetobacter* system can be explained by a natural selection process that does not require a stress response. These findings have fundamental implications to understanding the role of growth-limiting selective environments on cancer development.

## Introduction

Gene amplification mutations occur across the diversity of life, from viruses to humans (Romero and Palacios 1997; Gaines et al. 2010; Elde et al. 2012). They play important roles in both short-term adaptation to unfamiliar environments and long-term evolution of novel genes (Ohno 1970; Reams and Neidle 2004b; Andersson and Hughes 2009; Kondrashov 2012; Blount et al. 2012; Elliott et al. 2013). In recent years, amplification mutations have been established as key factors contributing to human cancers, in which gene amplification promotes tumorigenesis and chemotherapy resistance (Debatisse and Malfoy 2005; Albertson 2006; Santarius et al. 2010). In pathogens, amplification mutations contribute to increased virulence and antibiotic resistance (Craven and Neidle 2007; Sandegren and Andersson 2009). Still, the underlying role of selective environments in driving gene amplification formation remains unclear for any organism.

Amplification mutations initially arise in clustered (i.e. focal) tandem arrays, as either direct or inverted oriented repeats (fig 1) (Roth et al. 1996; Reams and Roth 2015). If an amplified genomic segment encompasses an entire gene(s), its expression is increased due to the increased copy number of the amplified gene’s promoter(s) (Iantorno et al. 2017). Interestingly, *de novo* mutants carrying high-copy amplifications (>2 tandem copies) encompassing entire genes have been reported only under selective conditions where the amplified gene’s increased expression provides a growth advantage (Wrande et al. 2008; Patterson et al. 2018). In two separate well-studied bacterial model systems, the number of nascent amplification mutant colonies continually increases over time during prolonged exposure to nonlethal selective minimum lactose agar medium (Cairns and Foster 1991; Quiñones-Soto and Roth 2011; Quiñones-Soto et al. 2012). The theory of adaptive amplification explains these observations by proposing amplification mutations are inducible by “selective stress” (Hastings et al. 2000). According to this theory, a proposed stress response that generates amplification mutations is stimulated under selective conditions due to its inherent growth inhibition on unadapted nonmutant cells. This theory has been implied to all organisms, including humans where gene amplification is found in cancer cells (Rosenberg et al. 2012). If true, adaptive amplification theory infers diverse organisms have evolved mutagenic stress responses to stimulate gene amplification mutations when cells are placed under growth-limiting selective conditions.

**Figure 1.**
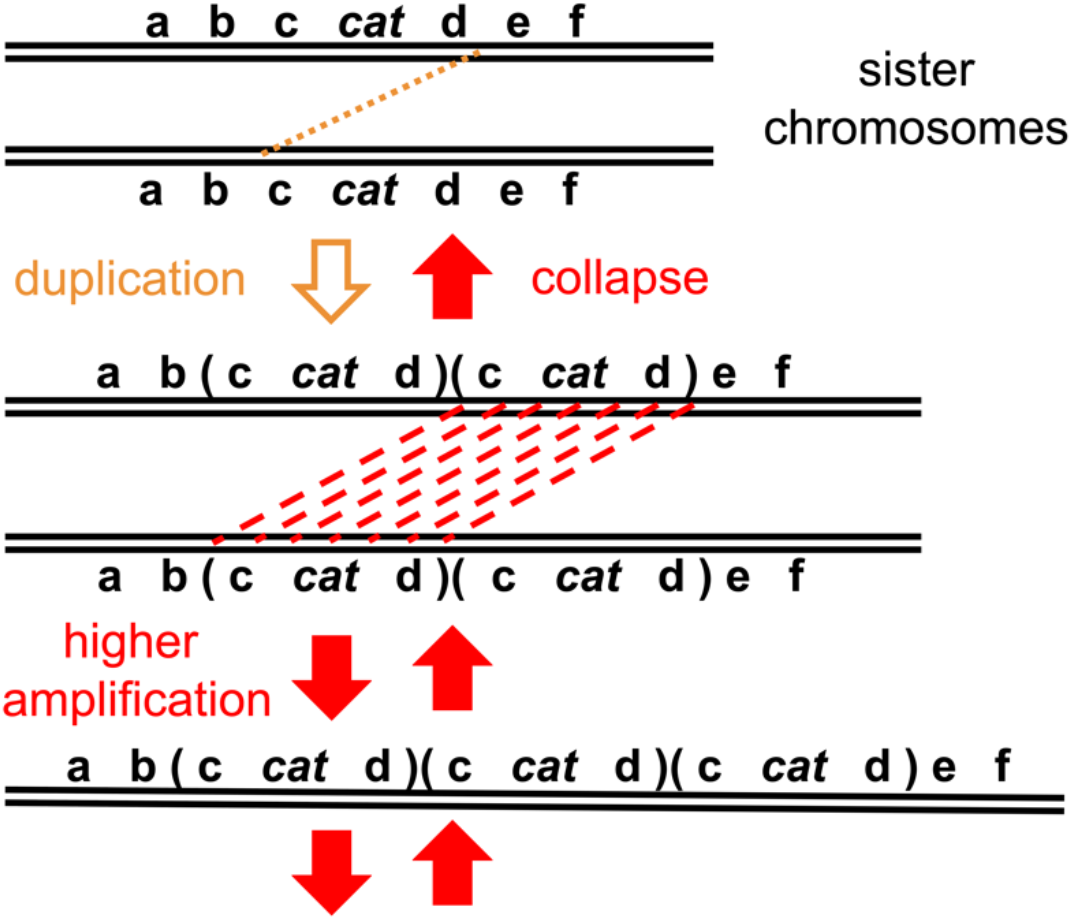
Tandem head-to-tail gene amplification mutations form by a two-step process. The top line represents the *cat* gene cluster region on the wild-type *A. baylyi* chromosome. In the first duplication step (yellow downward arrow), a cross-over event occurs between two distant loci (d and c) on sister chromosomes, indicated by the yellow dotted line. In the Ben reversion system, these duplication events predominantly depend on illegitimate recombination between regions of very short to no shared sequence identity (0-5 bp). The chromosomal region that is amplified (amplicon), shown by parentheses, vary widely in size (12-290 kb) between different Ben+ revertants, but always encompass the *cat* genes. During subsequent higher amplification steps (red downward arrows), the duplicated segments serve as extensive substrates for unequal crossing-over events between sister chromosomes, indicated by the red dashed lines, allowing them to occur at much higher rates than duplication formation. These secondary steps can occur multiple times in series, leading to incrementally higher amplicon copy numbers (e.g., 3, 5, 9, 17 copies, etc.). These same secondary interchromosomal recombination events, along with intrachromosomal exchanges, cause rapid collapse and loss of the amplification mutation (red upward arrows) once selection is removed.

Controversy surrounds this theory due to its implications to cancer and evolution, and whether selective environments influence the rates and actual processes of genetic variation. In contrast, an alternative theory, termed “amplification under selection”, argues amplification mutations arise without stress induction through a multistep clonal evolution process (Roth et al. 2006; Maisnier-Patin and Roth 2015). According to this modern Darwinian view, amplifications are initiated by partial variants (duplications) that arise during nonselective growth. Upon selection exposure, these partial mutants undergo multiple subsequent mutations (higher amplification) without any stress induction. Rather, each higher amplification step is gradually enriched over multiple cell divisions by their increased growth advantage through natural selection until a fully fit variant is achieved. Several researchers have argued the various cases apparently supporting adaptive mutagenesis are the result of artifacts in the experimental systems, and not due to evolved stress-induced mutagenic responses (Sniegowski and Lenski 1995). For these reasons the adaptive theories of evolved mutagenic mechanisms in nongrowing cells under selection are still heavily debated.

Here, we investigate both theories using a previously reported gene amplification model system in the bacterium *Acinetobacter baylyi* (Reams and Neidle 2003; 2004a; Seaton et al. 2012). Unlike other systems, this experimental system is distinctive by its ability to exclusively select for gene amplification mutants. This is due to a rigorous selection where growth on agar plates with benzoate as the sole carbon source requires mutations that increase expression of two distinct transcriptional units, the *catA* gene and the *catBCIJFD* operon. These *cat* genes, encoding enzymes for benzoate catabolism, are clustered within a 10-kb region on the circular chromosome. In the system’s parent strain, these genes have only low unregulated expression due to the absence of their two transcriptional activators. As a result, this parent strain is unable to grow on benzoate plates (Ben-). Only amplification of genomic segments encompassing this 10-kb clustered *cat* gene region singly increases the basal expression of all these *cat* genes to support colony growth on selective benzoate plates (Ben+) (fig 1). Since this selection yields only amplification mutants, it allows the process of gene amplification to be directly analyzed without the interference of other mutation types, such as frameshifts or point mutants.

Using this experimental system, here we test the two theories explaining the role of selective environments on gene amplification formation by reconstruction experiments, Lederberg replica plating assays, and Luria-Delbrück fluctuation tests (Luria and Delbrück 1943; Lederberg and Lederberg 1952; Hastings et al. 2000). The goal of these three independent methods is to determine whether the amplification mutations arise before or in response to selective stress. Based on our experimental results, we propose a model on how *cat* gene amplification mutations arise during selection. We discuss the implications of these results on understanding the development of cancer.

## Materials and Methods

### Bacterial growth conditions

Unless otherwise noted, bacteria were cultured at 30°C in Lysogeny (LB) medium or minimal medium (MM) with 10 mM succinate or 2 mM benzoate as the carbon source (Sambrook 1989; Singh et al. 2019).

### Bacterial strains, Ben reversion assay, and lawn growth measurements

*Acinetobacter* spontaneous Ben+ amplification mutants were derived from the Ben-parent strain ACN293 (Reams and Neidle 2003). This strain was subsequently renamed DR1023. For the Ben reversion assay, DR1023 parent strain cells were streaked out from a −80°C frozen stock onto nonselective agar plates containing either LB medium or 10 mM succinate MM, as indicated. After incubation at 30°C, single colonies were inoculated into tubes or baffled flasks containing between 2 to 100 ml of either liquid succinate MM or LB broth, while consistently maintaining the same liquid medium type as the streak plate medium. Cultures were grown to saturation overnight at 30°C with 220 rpm shaking. Cells from each fully-grown culture were pelleted and washed three times with 0.85% saline solution to remove any residual carbon source. Washed cells were concentrated 10-fold by resuspending in saline solution at 1/10th volume of the overnight liquid culture. A 100 μL sample was spread plated on a selective plate with 2 mM benzoate as the sole carbon source (Ben plate). Each selective Ben plate was spread with approximately the same number of parent strain cells (∼5 x 10^8^ cells) and incubated at either 30°C or 22°C, as indicated. The number of viable cells spread on each Ben plate was calculated by performing serial dilutions on the remaining concentrated cells, with CFUs determined on LB plates. Ben plates were analyzed every 1 to 3 days for new Ben+ revertant mutant colonies. The term “Ben+ revertant” refers to the return or “reversion” of the Ben-DR1023 parent strain back to the wild-type *A. baylyi* ADP1 strain’s Ben+ phenotype.

At various times points, the population of viable parental tester cells in the lawn on the Ben plates was determined by taking agar plugs from the surface of the selection plates, avoiding any visible Ben+ colonies. Cells from the plugs were suspended in 0.85% saline solution, vortex mixed for at least 2 h, and serially diluted. Dilutions were plated on LB plates and used to calculate the lawn viable cell count on the selective plates.

### Preparation of intact genomic DNA, restriction digestion, pulsed-field gel electrophoresis (PFGE) analysis, and Southern hybridization methods

DNA samples for PFGE analysis were prepared in agarose plugs as described in Gralton et al. 1997 (Gralton et al. 1997). Plugs were digested with the restriction enzyme NotI and electrophoretically separated by the pulsed-field methods of either transverse alternating field electrophoresis (TAFE, Geneline II, Beckman) as previously described (Gralton et al. 1997), or CHEF-DRII using a 24 h protocol with a constant 6-V cm^-2^ current and pulse switch times that increase from 10 to 200 s. Southern hybridization methods were carried out as indicated by Gralton et al. 1997 (Gralton et al. 1997). Nonradioactive probes, prepared with digoxigenin and a random-primed labeling system (Genius Systems, Boehringer Mannheim), were hybridized to target sequences, and detected with anti-digoxigenin alkaline phosphatase conjugates and chemiluminescent substrates. Hybridization signal intensities on exposed X-ray films were digitized and calibrated on a UVP Lab Products gel documentation system according to the directions of the LabWorks image acquisition and analysis software.

### Measurement of amplicon length and copy number

For all amplification mutants, amplicon length and copy number were estimated by PFGE and Southern hybridization as previously described (Reams and Neidle 2004a). The amplicon size for 74 amplification mutants was verified by isolating and sequencing over their junction endpoints (Reams and Neidle 2003). The distance between them was calculated based on the *A. baylyi* ADP1 genome sequence. This calculation accounts for the 446 bp *catM* deletion in strain DR1023.

### Reconstruction experiments

Reconstruction experiments were performed by plating a known number (approximately 300) of Ben+ revertant cells with about 5 x 10^8^ DR1023 cells as in the original Ben reversion amplification mutant selection. Multiple Ben+ revertants arising on various days were isolated and tested. Additional assays were conducted with derivative ACN277 strain cells carrying different amplicon copy numbers (15, 8, 3, or 2). Before plating, each population of cells was grown in 10 mM succinate minimal medium. Background amplification rate was determined by plating 5 x 10^8^ DR1023 cells without any Ben+ revertant cells onto selective plates. Each set of experiments with different Ben+ revertants arising on various days were repeated three times, while experiments with ACN277 and its derivative strains were repeated ten times.

### Replica plating experiments

LB cultures of the Ben-DR1023 parent strain were grown overnight to saturation, serially diluted, and 5,000 to 50,000 CFUs were spread plated onto multiple rich medium (LB) agar plates and incubated at 30°C. After 13-16 days incubation, each LB plate was replicaprinted onto one LB plate followed by three selective Ben plates. The position of each replica plate was marked to determine whether Ben+ revertants arise in the same positions. The nonselective LB plates were incubated for 2-3 days then stored at 4°C for future analysis. The sets of three selective plates were incubated at 30°C for up to 21 days and analyzed every 1 to 3 days for new Ben+ revertant colonies. Any clustered Ben+ revertant colonies arising in the same location on all three selective plates were recorded as potential siblings and separately grown in liquid benzoate MM for further analysis.

To isolate and purify in the absence of selection the progenitor (precursor) cells that gave rise to the selected Ben+ revertants, cells were collected from the LB replica plate at the corresponding location to the revertant colonies on the three Ben plates. These cells were grown overnight to saturation in LB liquid medium at 30°C. Cultures were diluted and under 500 CFUs were spread plated onto LB agar plates. After incubation at 30°C for 2-3 days, these plates were once again replicaprinted onto one LB plate followed by three Ben plates. The LB plates were incubated for 2-3 days then placed at 4°C for future analysis. The sets of three Ben plates were incubated at 30°C for up to 14 days and analyzed every 1 to 3 days for new Ben+ revertant colonies. Successful enrichment of precursor cells was indicated by many same-position clustered Ben+ colonies on all three Ben plates. Clonal populations of precursor and revertant cells were collected and saved from isolated colonies at the same position on the replica plates.

### Fluctuation tests

122 independent parallel cultures of the Ben-DR1023 parent strain were grown overnight in nonselective LB media and were then tested for Ben+ mutant frequency using the Ben reversion assay. Each culture was derived from a different single isolated colony. For 3 of the 122 cultures, a flask was used to grow large cultures (40-100 ml). The other 119 cultures were grown in smaller volumes (2 ml each). The cells from the large cultures were plated onto 18-49 selective plates for each culture, whereas the smaller culture cells were plated onto one selective plate per culture. Regardless of the culture volume, each selective plate was spread with approximately the same number of parent strain cells (5 x 10^8^ cells). Selective plates were incubated at 30°C and analyzed every day for 21 days. The distributions of mutant frequencies were analyzed by plotting the relation between the logarithm of the proportion of selective plates with *x* or more Ben+ mutants (y-axis) and log *x* (x-axis) (Luria and Delbrück 1943; Cairns et al. 1988; Cairns and Foster 1991).

### Statistical Analysis

Correlation between amplicon length and day number mutant colony arose was assessed with both Spearman Rho and Kendall Tau correlation tests. The Brown-Forsythe equality of variance test was performed to determine whether there was a significant difference in variances of Ben+ revertant frequencies per Ben plate measured between multiple cultures versus within single cultures (Brown and Forsythe 1974a; 1974b). All three statistical tests were independently performed in both R and Excel 2016 to assure consistent results.

## Results

### The emergence of Ben+ amplification mutant colonies increases exponentially during prolonged selection on nutrient-limiting plates

In the Ben reversion assay, spontaneous Ben+ mutant colonies are selected by placing a population of the Ben-parental cells, *A. baylyi* strain DR1023, onto minimum medium agar plates with benzoate as the sole carbon source (Ben plates). During prolonged incubation on the selective Ben plates, Ben+ mutant colonies accumulate exponentially over time (fig 2). Rare Ben+ revertant colonies are visible as early as day 3 while additional new visible mutant colonies continually arise and accumulate over time until the plates are either saturated with revertant colonies or dried out after about 50 days. Our previous studies have shown these amplification mutant colonies accumulate to a frequency of 10^-8^ (Ben+ amplification mutant colonies per total cells plated) within 21 days (Reams and Neidle 2003). Here, with the aim of making this assay more efficient, we compared the effect on reversion frequencies of varying the growth media before selection (minimum succinate vs rich LB) and the selective incubation temperatures (22° vs 30°C). Under all conditions tested, the number of Ben+ revertant colonies that were selected on the Ben plates increased exponentially over time. Although the growth medium before selection did not significantly affect the mutant frequencies, selection at 30°C compared to 22°C caused an approximate 10-fold increase in revertant accumulation (fig 2A). Under these optimized conditions, Ben+ amplification mutant colonies accumulate to frequencies of 10^-9^ at 9 days, 10^-8^ at 14 days, and 10^-7^ at 19 days (fig 2B). To confirm Ben+ colonies form exclusively by gene amplification under these optimized conditions, 75 colonies appearing on various days were purified and analyzed by Pulsed-Field Gel Electrophoresis. All had increased gene dosage of the *cat* chromosomal region. Moreover, no Ben+ colonies arose within two days of incubation.

**Figure 2.**
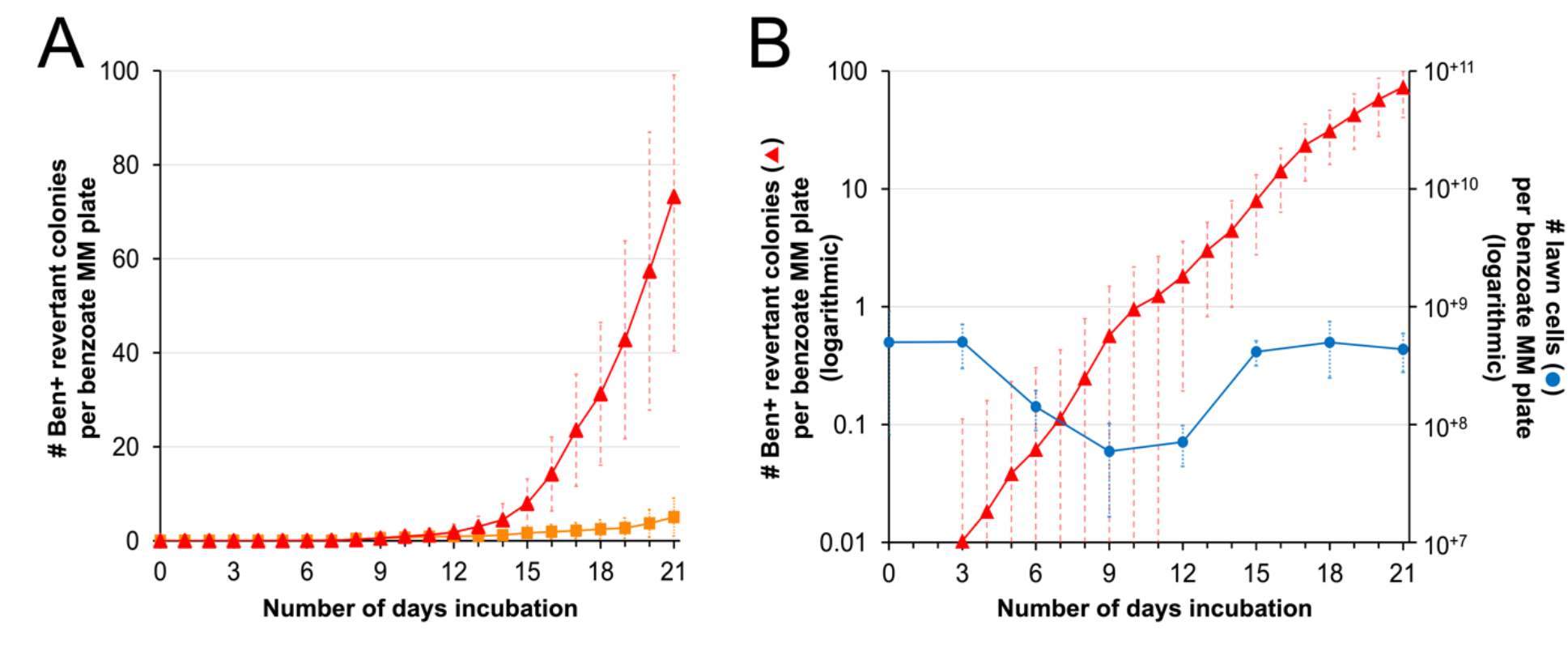
Ben+ revertant colonies carrying nascent *cat* gene amplification mutations accumulate exponentially over time on selective Ben plates. A) Comparing Ben+ revertant frequencies using previous assay conditions (succinate MM growth before selection, selective incubation temperature at 22°C) indicated by orange square versus new optimized conditions (LB growth before selection, selective incubation temperature at 30°C) shown by red triangles. Each point represents the average value from 100 independent repeated trials. This data was derived from 119 trials where 19 outliers (jackpots) were removed. Error bars represent the standard deviations across the 100 independent trials. B) Ben+ revertants accumulate exponentially both under the new optimized (red triangles) and previous test conditions (not shown). The Ben+ revertant frequencies represent the same data as figure 2A but graphed with the left y-axis on a logarithmic scale. Changes in the lawn cell population size (blue circles) are indicated by the right y-axis. Both y-axes are graphed proportionally to each other on a logarithmic scale to allow a direct comparison between lawn population size changes and Ben+ revertant accumulation.

### Investigating potential causes for the delay in Ben+ amplification mutant colony formation

Most Ben+ amplification mutant colonies do not become visible until after a significant delay. For example, the majority of accumulated revertant colonies over a 21-day period do not become visible until after 18 days of incubation (fig 2). This delay in colony formation was not a direct effect of the slow growth of the amplified mutants since when cells were transferred from their original selective plate to a fresh selective plate they formed large single colonies similar to the growth rate of the wild type, within 1-3 days. We also tested whether the diverse times in Ben+ colony formation under the original selection could be explained by a correlation between the day the colonies appeared and the size of the amplified chromosomal *cat* gene region (amplicon) (fig 3). However, no practical significance in the correlation was detected using both Spearman and Kendall correlation tests for data collected during 30-days incubation (Spearman coefficient of 0.27 with a p-value of 0.016 and Kendall Tau coefficient of 0.20 with a p-value of 0.012). To analyze where the strongest correlation exists, the data was subdivided into three groups with 10-day ranges. Using a Kendall Tau correlation test, only the first period (days 3-13) indicated a very weak correlation between the time until colony appearance and amplicon size (Kendall Tau correlation coefficient of 0.38 with a p-value of 0.017). On one hand, only revertants with smaller amplicons (<100 kb) arose prior to 8 days, whereas those with larger amplicons (>100 kb) required a minimum of 8 days of incubation time. On the other hand, after 11 days of incubation, mutants with wide varieties of amplicon sizes formed visible colonies after the same length of incubation. For example, 3 independent amplification mutants arising on the 11^th^ day had amplicon sizes of approximately 32, 146, and 273 kb, respectively. Therefore, in general, amplicon size alone does not fully explain the delay in colony formation in the Ben reversion system.

**Figure 3.**
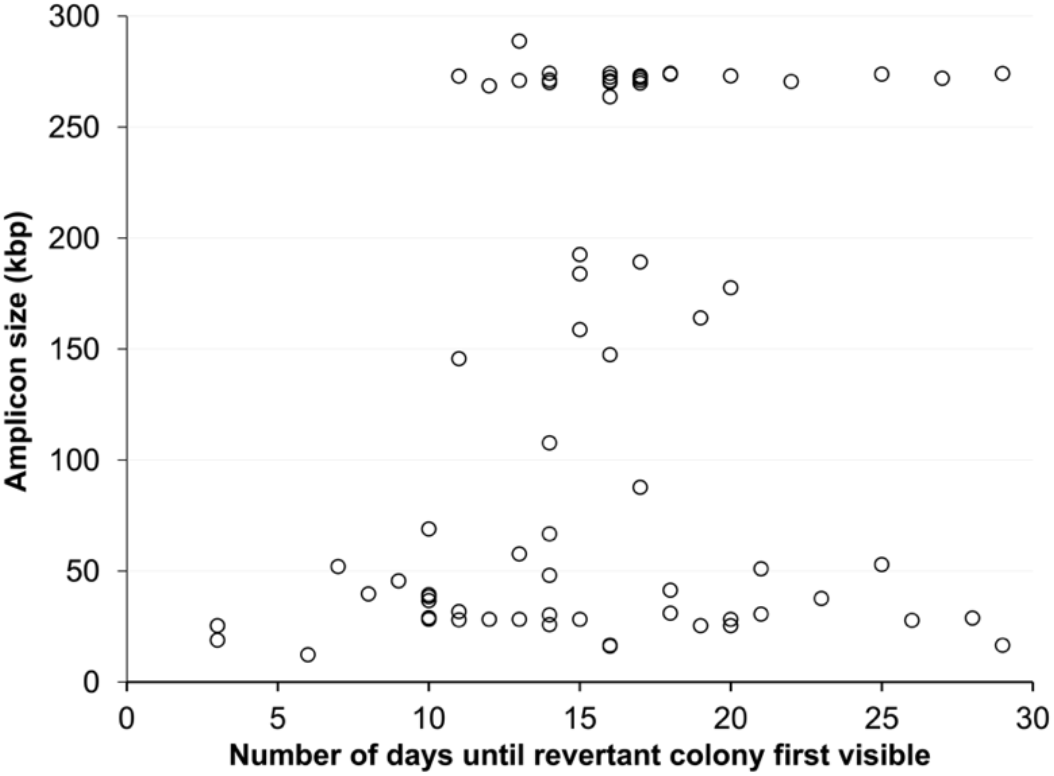
Time until visible Ben+ colony formation does not directly correlate with amplicon length. Amplicon size (y-axis), determined by PFGE and join point sequence analysis, for 74 independently isolated Ben+ revertants that were first visible after different days of incubation (x-axis) where the Ben plates were incubated for 30 days. A Spearman Rho test yielded a correlation coefficient of r_s_ = 0.27 with a p-value = 0.016. A Kendall Tau test produced similar results with a correlation coefficient of r*_τ_* = 0.20 with a p-value = 0.012.

Additionally, we investigated whether the exponential accumulation in revertant colonies during prolonged incubation could result from residual growth of the Ben-lawn population. At various time points, viable cells on the surface of the Ben plates were quantified by taking agar plugs of the lawn cells, avoiding any visible Ben+ revertant colonies, and CFUs from these plugs were assessed. While the Ben+ revertant frequency increased exponentially nearly 10,000-fold between 3 to 21 days, the lawn population change was negligible (fig 2B). Initially, the lawn population size slightly decreased (∼10-fold) during the first 12 days of incubation, followed by a minor increase to the original starting population size by day 21. Therefore, the lawn cell population does not match the exponential increase in Ben+ revertant frequency during extended selection.

### Do high-copy cat gene amplification mutants arise before or during exposure to growth-restrictive selective conditions?

The delay in Ben+ revertant colony formation could be explained by the amplification mutations forming on the selective Ben plates. To test whether the Ben+ amplification mutations occurred before or during exposure to selection, we performed reconstruction experiments as previously used in studies investigating the adaptive amplification theory (Hastings et al. 2000). The purpose of these experiments is to understand why selected Ben+ colonies appear late and after a delay, even though they form colonies rapidly (1-3 days) when transferred to fresh selective plates. If fast-growing Ben+ mutants are present in an initial population prior to selective nutrient starvation on Ben plates, the delayed appearance of colonies might be attributed to competition amongst the high density of parent strain cells (5 x 10^8^ CFUs/Ben plate). This competition may be growth restricting and cause the delay in colony formation in the Ben reversion assay. Therefore, transferring a Ben+ colony to fresh selective medium does not represent the same conditions as the original selection. To assess the time needed for Ben+ amplification mutants to form visible colonies under the original selective conditions, Ben+ revertant cells carrying high-copy amplification were diluted and plated along with a high density of Ben-parent strain cells onto selective medium. These conditions mimic and “reconstruct” the original selective conditions in which the amplification mutants initially arose.

Reconstruction experiments were conducted with 25 independent Ben+ revertants that initially arose after a range of incubation periods, from 4-21 days. In all cases, revertants formed visible colonies more quickly in the reconstruction experiment than in its corresponding original isolation (fig 4). For example, a revertant strain that appeared as a colony on day 14 in the original selection formed colonies within only 3 days in the reconstruction experiments. On average, Ben+ colonies arose 8 days faster in reconstruction experiments compared to their original selection. Therefore, the presence of a high density of parent strain cells did not cause rare high amplification cells to grow slowly. These results suggest cells with high-copy gene amplification did not exist in the starting population and that some aspect(s) of the gene duplication and higher amplification process occurred during the extended period of selection.

**Figure 4.**
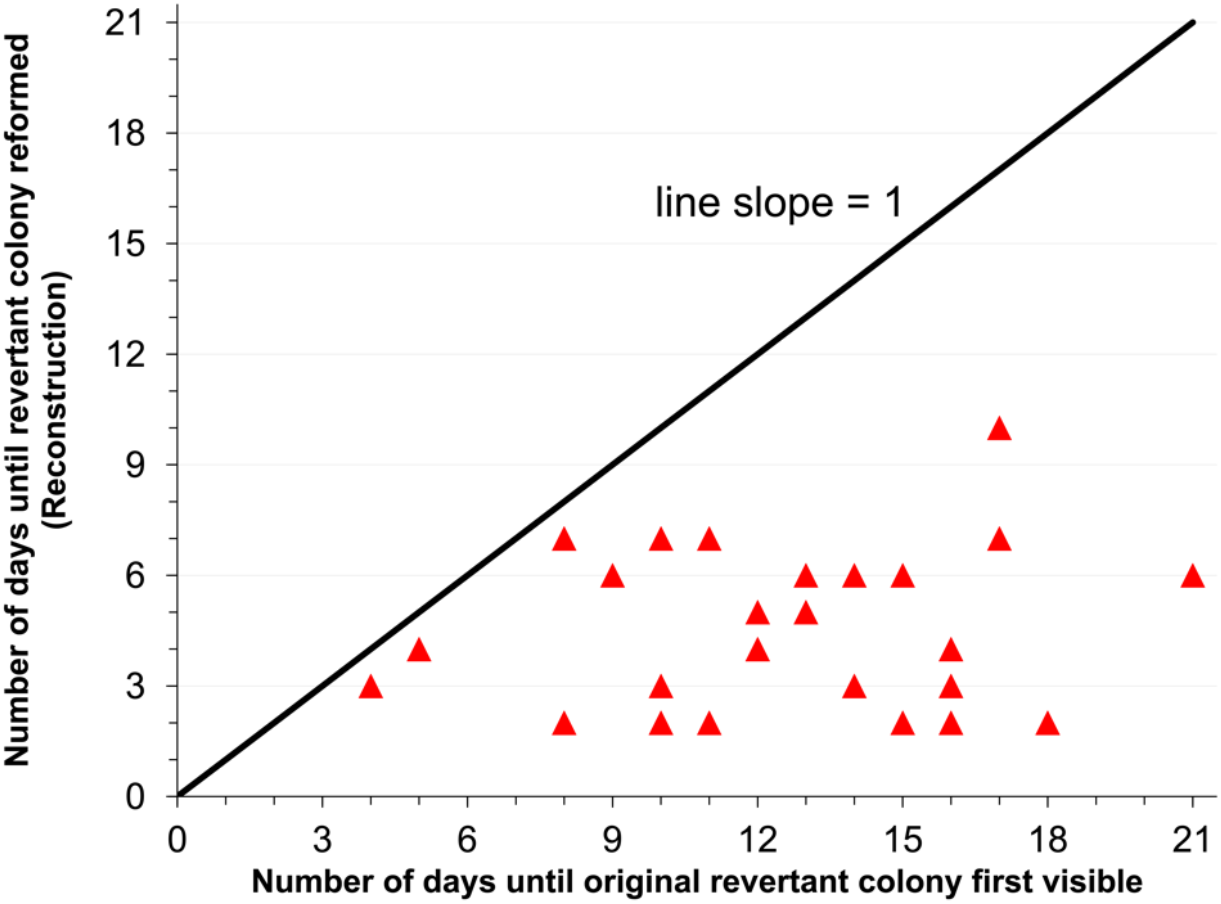
Reconstruction experiments with high-copy amplification mutants. Each red triangle represents an independently isolated Ben+ revertant verified to carry a high-copy *cat* gene amplification mutation by PFGE. The x-axis shows the number of days it took for the revertant to form a visible colony under the original isolation. The y-axis indicates the number of days it took the revertant to reform a visible colony under the reconstruction conditions, which mimic the original selective conditions. The diagonal line with a slope of 1 serves as a reference to determine whether the revertant took the same (along line) or less (below line) time to form a visible colony under the reconstruction conditions. Reconstruction experiments were repeated three times for each revertant; in all 25 revertants, standard deviations were within one day between the three trials.

### Ben+ colony formation with strains carrying different levels of cat gene copy numbers

The results of the reconstruction assays suggest fast-growing Ben+ cells with high *cat*-region gene dosage do not exist in the initial population from which revertants were selected. To explain the emergence of Ben+ colonies, the adaptive amplification theory posits that gene duplications, as the rate-limiting step in the formation of amplification mutants, would be induced during exposure to growth-restrictive selective conditions. An alternative possibility proposed by the amplification under selection theory is that cells within the initial population carry *cat* gene duplications or other low levels of gene amplification. These variant cells could give rise to Ben+ revertant colonies as higher *cat* gene amplification is selected during growth on benzoate. To understand the appearance of Ben+ colonies from cells with different initial levels of *cat* gene amplification, we isolated several derivatives of a Ben+ amplification mutant. Each derivative population contained cells with a reduced copy number of the *cat* gene region.

Using a previously described method (Reams and Neidle 2003), a Ben+ revertant strain, ACN277, carrying high-level tandem amplification (15 copies) of an approximately 28 kb amplicon was grown under nonselective conditions with succinate as its carbon source. Such growth conditions do not require amplification for growth and continual passage of ACN277 resulted in derivative strains with fewer amplicon copies (fig 1). After 200 generations of nonselective growth, derivative strain populations were isolated in which the majority of cells carried different copy numbers of the 28 kb *cat* gene amplicon. A frozen stock of each derivative cell population was made by growing individually isolated colonies on succinate to obtain a sufficient number of cells for DNA isolation and analysis. This procedure corresponded to approximately 30 generations of growth on nonselective medium. To determine the amplicon copy number of each derivative population, plugs were prepared and analyzed by PFGE and Southern Hybridization.

Figure 5B shows the results for three of these derivatives that varied in amplicon copy number. Each of these three derivative populations appears to represent different intermediate stages in the process of deamplification. Derivative populations were obtained in which most cells contained 2, 3, and 8 copies of the 28 kb amplicon. These populations were designated 2X, 3X, and 8X, respectively.

**Figure 5.**
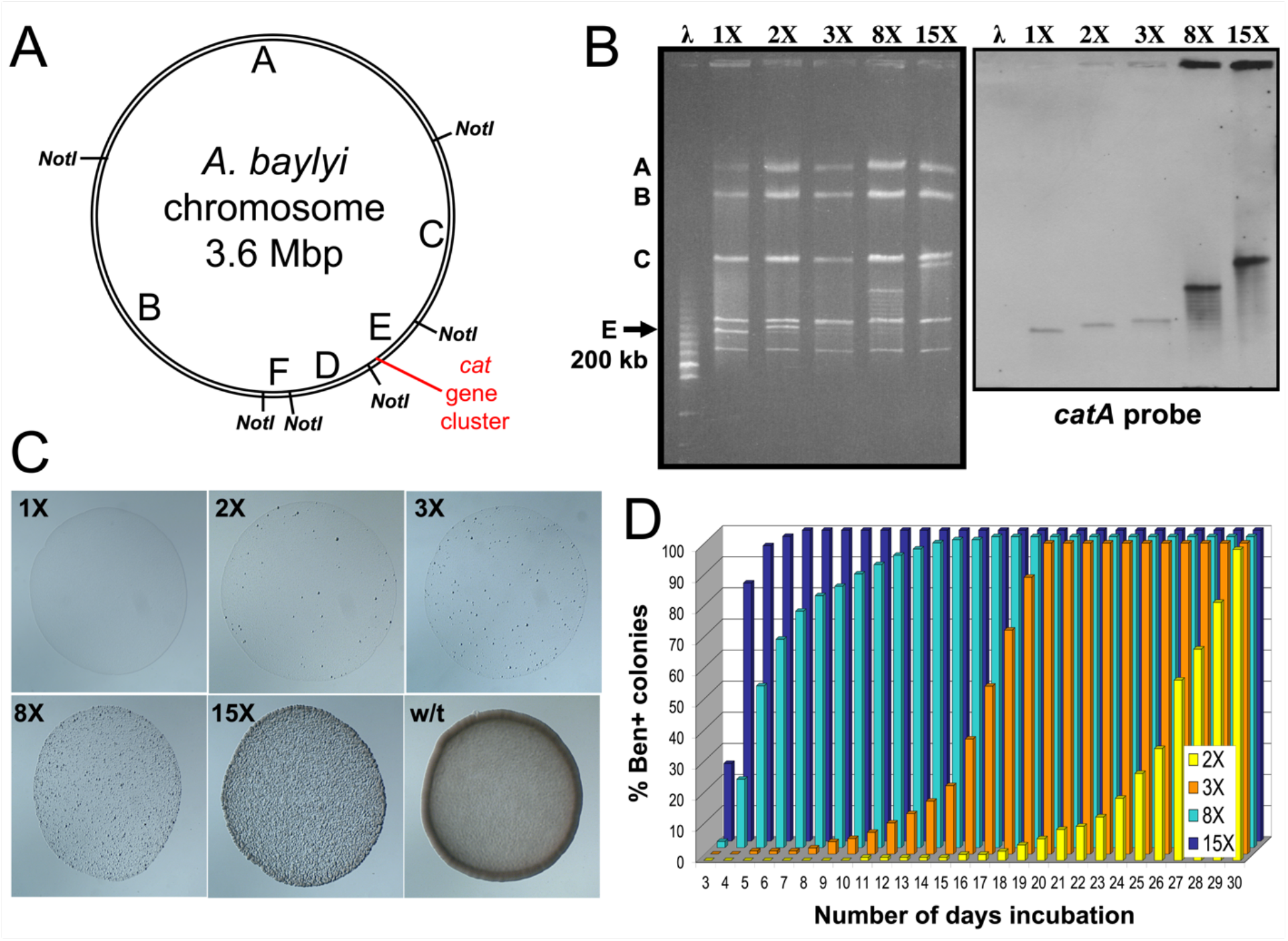
Reconstruction experiments with strains carrying different *cat* gene copy numbers. A) Chromosomal map of *Acinetobacter baylyi* showing the locations of NotI cut sites and the *cat* gene cluster region. NotI digestion of the wild-type chromosome yields six fragments designated A to F in decreasing size order: A 1,380 kb, B 1,044 kb, C 622 kb, D 259 kb, E 197 kb, and F 97 kb. The *cat* genes are located on the normally ∼200 kb E fragment. B) PFGE (left) and Southern hybridization (right) analysis of strains carrying different copy numbers of a ∼28 kb amplicon encompassing the *cat* gene region. Genomic DNA of each strain was digested with NotI. A *catA* probe was used for the Southern hybridization. The Lambda DNA size ladder (λ) was used as a size marker. The A, B, C, and E labels mark their corresponding NotI fragment size in the wild type. 1X refers to the haploid DR1023 parent strain, while the 2X, 3X, 8X, and 15X strains are strains carrying 2, 3, 8, and 15 amplicon copies, respectively. C) Phenotypes of strains carrying different *cat* gene copy numbers on selective Ben plates. The wild-type ADP1 strain (w/t), which does not require gene amplification mutations to grow on benzoate medium, is shown as a comparative reference. Each strain was grown to saturation in nonselective succinate liquid medium. After overnight growth, 5 μL of each undiluted culture was drop plated at different positions on a single Ben plate. Each image is a microscopic view of the corresponding strains’ 5 μL drops after 2 days incubation at room temperature (22°C). D) Reconstruction experiments with strains carrying different levels of *cat* gene amplification. Approximately 300 cells of each derivative strain along with 5 x 10^8^ cells of the DR1023 parent strain were spread plated onto each Ben plate. The percentage of Ben+ colonies refers to the total number of Ben+ colonies above background colonies divided by 300 colonies. The background colonies were measured by plating 5 x 10^8^ cells of the DR1023 parent strain without the derivative strains and counting the number of Ben+ revertant colonies each day. Each point represents the average value from 10 independent repeated trials with 10 plates per trial.

Each strain carrying partial amplification was grown overnight in nonselective succinate medium. After growth to saturation, 5 μL was dropped onto Ben plates and incubated at 22°C for 2 days (fig 5C). In general, the lower the average copy number, the longer the population took to grow when transferred back to selective Ben plates. Further analysis by PFGE and Southern hybridization of these benzoate-grown derivatives demonstrated they had regenerated higher amplification during selective growth. In contrast, strains which reverted completely back to a haploid state (1X) were unable to grow on benzoate medium, nor did they generate higher amplification.

Reconstruction experiments were performed with the high and intermediate level amplification strains (fig 5D). Where the high amplification revertant colony initially became visible after 12 days on the original selective medium, most of the cells carrying the preexisting high amplification (15 copies) arose within only 4 days under the conditions of the reconstruction experiments. Based on these results, higher amplification was generated under the original selective conditions and was not preexisting before selection.

In the reconstruction experiments with the intermediate copy number strains, the results were significantly different (fig 5D). The lower the initial copy number, the longer the cells took to form visible colonies. For example, most cells with only two copies took at least 27 days to form a visible colony under the original selective conditions. PCR analysis was performed on chromosomal DNA from 30 Ben+ colonies that arose after 20 days of incubation in these reconstruction experiments with the two-copy derivative. Of these 30 colonies, 12 contained the same duplication junction present in the 2X derivative. The other 18 colonies were considered spontaneous Ben+ mutants derived from the dense population of parent strain cells. PFGE and Southern hybridization analysis on these 12 strains showed a ladder of bands hybridizing to a *catA* probe indicating that all colonies had regenerated higher amplification from the preexisting duplication.

In contrast to the 2X population, where most cells took at least 27 days to form a visible colony, most cells in the 3X and 8X populations took at least 17 and 5 days to form a visible colony, respectively (fig 5D). Since these low copy number strains generate higher amplification when grown on benzoate, these reconstruction experiments indicate the secondary steps of generating higher amplification required several days to occur, that were increasingly longer with lower initial copy numbers. These results suggest these secondary steps are the rate-limiting event to Ben+ colony formation. They further suggest the generation of higher amplification from a low-level amplification, such as a *cat* gene duplication, is the cause for the delay in visible colony formation under the original selective conditions.

### Ben+ gene amplification mutants originate before selective pressure

Our experiments indicate Ben+ high-copy amplification mutants are not present before exposure to the selective starvation conditions, however it is still possible low-copy cells are present in the initial unselected population that are predecessors to high-copy Ben+ revertants. To assay whether these ancestral cells are present before selective pressure, replica plating experiments were performed as described previously (Lederberg and Lederberg 1952). The goal of these experiments is to detect and potentially isolate predecessor cells that give rise to the selected Ben+ revertant colonies before the cells are exposed to selection. For these experiments, approximately 5,000 to 50,000 microcolonies of the Ben-parent strain were cultivated on individual nonselective (LB) plates, where growth does not require gene amplification (fig 6A). These plates were replicaprinted onto four plates: one LB plate (providing a library of all cells) and three selective Ben plates (where amplification mutations are required for growth). The position of each replica plate was marked to allow alignment and determine whether Ben+ revertants arise in the same positions.

**Figure 6.**
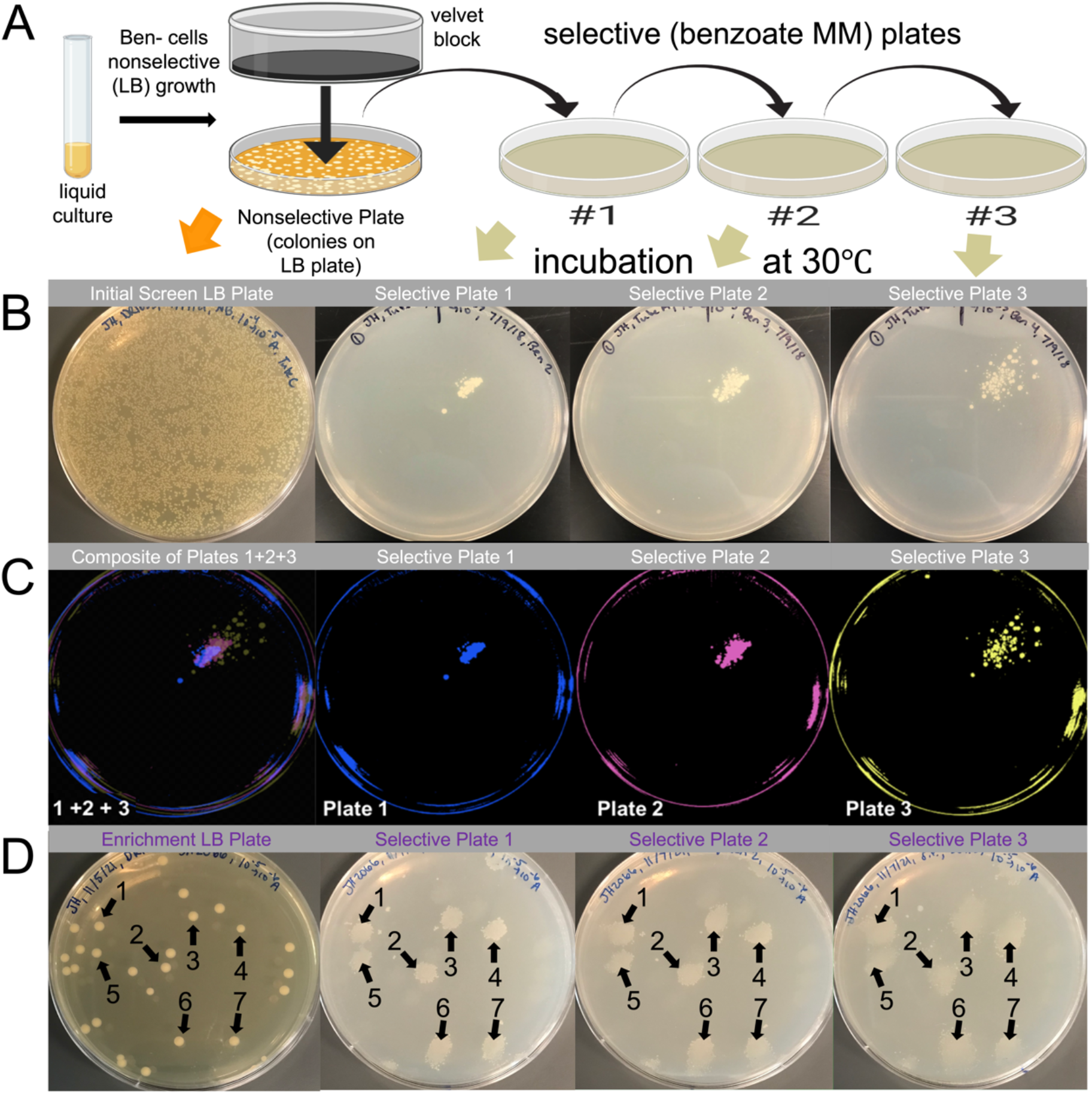
Replica plating assays to detect unselected progenitor cells of Ben+ revertants. A) Schematic of replica plating procedure: a nonselective liquid culture of the Ben-parental strain is diluted and plated onto a nonselective plate. After incubation, colonies are replicaprinted onto three selective Ben plates, incubated at 30°C, and monitored for growth. Image created with BioRender.com. B) A nonselective plate and its three selective replica plates showing clustered Ben+ revertant colonies arise in the same position on all three selective plates. C) The selective plates 1, 2, and 3 in panel B were processed using GIMP software to fluorescently highlight the Ben+ colonies blue, pink, and yellow, respectively. The plate on the far left is an overlap composite of these three fluorescent plates, where the overlap of the three colors yields a new color: purple. This purple confirms the colonies were in the same position and was used to locate the corresponding position on the nonselective plate to enrich the ancestral precursor cells. D) Enriched precursor cells were collected from the nonselective plate, grown in LB, diluted, and spread onto a fresh nonselective plate to repeat the replica plating procedure in panel A. Like panel B, this panel shows a nonselective plate (with isolated colonies) and its three selective replica plates. The numbers 1-7 indicate clustered Ben+ colonies present in the same position on all three selective plates and the nonselective plate.

Our replica plate experiment results show single Ben+ revertant mutant colonies accumulate on the printed selective Ben plates during prolonged incubation, consistent with the standard Ben reversion assay. However, in contrast to the standard assay, additional clustered Ben+ revertant colonies arose on the printed selective plates (fig 6BC). These clustered colonies appeared as multiple colonies congregated within one position on the selective plate. Typically, these clustered colonies were the first revertants to appear on the selective plates, becoming visible after 3 to 14 days of incubation, but appeared on later days as well. Importantly, all clustered revertant colonies arose in the same position on the three selective replica plates. We performed these replica plate experiments for over 100 independent DR1023 cultures, and repeatedly observed these same-position clustered revertant colonies across different cultures. These same-position clustered colonies appeared as though a revertant colony on the nonselective plate had been printed to the selective plates. Consistent with this possibility was the clustered colonies became more diffuse with each printing such that the clustered Ben+ revertants were more smudged on the third and final selective plate. These results suggest these mutants were present on the nonselective plate and therefore formed before selective stress was imposed.

To isolate the progenitor cells that gave rise to the Ben+ revertants on the three selective replica plates, cells within the corresponding position on the nonselective plates were collected, regrown in LB liquid overnight, and further analyzed by repeating the replica plating assay. During this enrichment process, replica plating was performed with nonselective plates carrying fewer colonies (<500 colonies per plate) than the initial round (>5,000 colonies per plate). After replica plating, we consistently observed a high frequency of same-position clustered revertants on all three selective replica plates. An example of these enrichment replica plating results is shown in figure 6D. In repeating these tests, 10% to 90% of the colonies on the nonselective plate gave rise to same-position clustered Ben+ revertants. This wide variation in enrichment was most likely due to the difficulty of precisely identifying the corresponding position and inadvertently picking up neighboring cells from the nonselective plate, where microcolonies are densely packed and not spatially isolated. Also, if ancestral precursor cells carry low-copy amplification, they are likely very unstable due to the absence of selection and collapse back to parental haploid during their nonselective growth. As a control, cells were collected from random positions of the initial screen nonselective plate that did not correspond to clusters of Ben+ colonies. In these cases, the results were identical to the initial round of replica plating. That is, very rare to no same-position clustered revertants were observed on the selective replica plates. Collectively, these results strongly suggest Ben+ revertants carrying high *cat* gene dosage arise from cells existing in the population prior to exposure to selective starvation. Importantly, they demonstrate the original ancestral cells can be enriched without the cells ever being exposed to selective stress.

### Ben+ revertants with high-copy cat gene amplification derive from cells with duplications or low-level amplification that exist before selective stress

To characterize chromosomal rearrangements in isolates from these replica plate experiments, such as the one shown in Figure 6D, we performed PFGE on genomic DNA digested with NotI (fig 7). Cells were analyzed from a well isolated colony that was never exposed to selective starvation (labelled at “Pre”), and the Ben+ revertants (labelled as “Rev”) that arose in the same position on the three replica plates (labelled A, B, and C). This method was repeated for three independent sets (labelled 1, 2, and 3). As a reference, a parent strain control, labelled “Ben-”, was used to indicate the NotI digest pattern in an isolated colony that did not give rise to Ben+ growth on selective medium.

**Figure 7.**
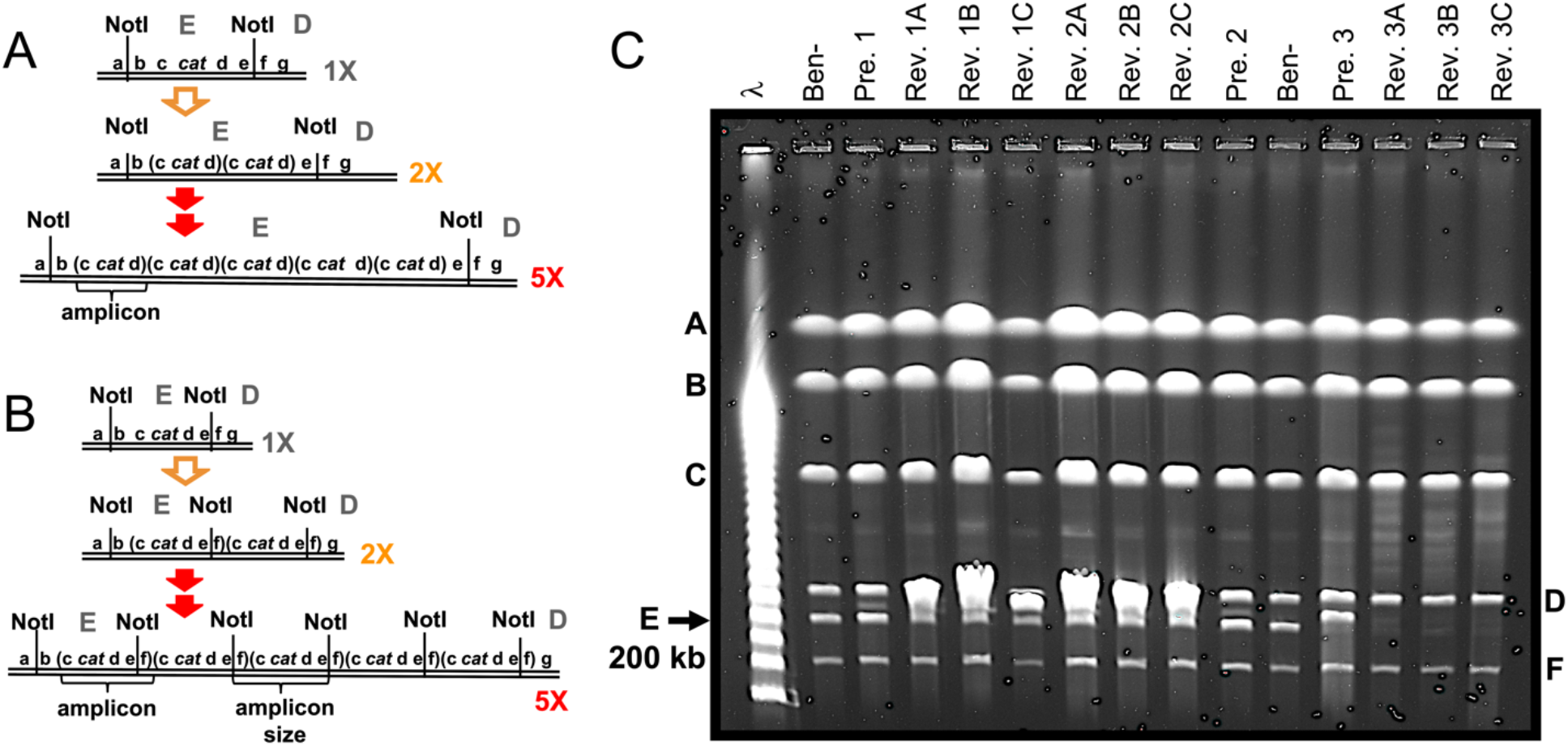
PFGE analysis of unselected precursors (Pre.) and Ben+ revertants (Rev.) from replica plates. A and B) NotI restriction maps of the *cat* gene region targeted for amplification in Ben+ revertants. A large-scale view of this map is shown in figure 5A. 1X refers to the parent haploid strain, while 2X and 5X refer to isolates carrying 2 and 5 amplicon copies, respectively. Panel A displays the scenario where the amplicon is confined within the NotI E fragment, causing an enlarged E fragment. Panel B shows the outcome where the amplicon includes the NotI cut site between the NotI E and D fragments (between e and f). This maintains the 200 kb E fragment but generates a new NotI fragment that corresponds to the amplicon size. C) PFGE analysis on three sets of precursors and their three Ben+ revertants. Each set has its own number. For example, the three revertants aligned with Pre. 1 are Rev. 1A, 1B, and 1C, where each revertant was from a separate replica plate. Ben-refers to the DR1023 parent strain. Genomic DNA from each isolate was prepared and digested with NotI. The Lambda DNA size ladder (λ) was used as a size marker. The A-F labels mark their corresponding NotI fragment size in the wild type: A 1,380 kb, B 1,044 kb, C 622 kb, D 259 kb, E 197 kb, and F 97 kb. The *cat* genes are located on the normally ∼200 kb E fragment.

Gene amplification of the *cat* gene region can be demonstrated on PFGE by two general NotI band patterns (fig 7AB) (Reams and Neidle 2003; 2004a). In revertants where the *cat* gene amplified segment is confined within the NotI E fragment, *cat* amplification is demonstrated by an enlarged E fragment, which houses the *cat* genes (fig 7A). This is the case for Revertants 3ABC which each have a missing 200 kb E fragment and several new larger faint bands, indicating enlarged E fragments, of different sizes between the D and C fragments (fig 7C). These multiple bands indicate a cell population that varies in amplicon copy number, where each faint band represents a subpopulation of cells carrying the same amplicon copy number.

In other revertants where the *cat* gene amplicon extends into the neighboring D fragment, the 200 kb E fragment is maintained but high *cat* amplification is demonstrated by an additional more intense NotI fragment (fig 7B). Since each amplicon contains a single NotI cut site and is thereby linearized by NotI digestion, this new band corresponds to the amplicon size and its relative higher intensity corresponds to the amplicon’s higher copy number. This is the case for Revertants 1ABC and 2ABC which each have an additional fragment between the D and E fragments of higher intensity (fig 7C). Estimated amplicon copy number in the 9 revertants ranged from 5 to 15 copies per cell, consistent with Ben+ revertants characterized under the standard Ben reversion conditions. Therefore, our PFGE results indicate all nine revertants carried high *cat* gene amplification.

Interestingly, the amplicon size was consistent across all three revertants within each set (fig 7C). The one set of revertants having enlarged NotI E fragments (Rev. 3ABC) yielded similar PFGE results across all three revertants isolated on separate selective replica plates. Invariably, these three revertants each contained a ladder of faint enlarged E fragments where each rung was separated by approximately 40 kb, indicating various subpopulations were present with different copy numbers of the 40 kb amplicon. Similarly, the other two sets of revertants had the same approximate amplicon size for all three revertants within its set: the Rev. 1ABC set had a ∼237 kb amplicon and the Rev. 2ABC set had a ∼247 kb amplicon. Together, these results support that each of these three sets of revertants are indeed siblings derived from a common ancestor present before selection.

In contrast to the high-copy *cat* gene amplification siblings, our PFGE results show the three ancestral precursor populations, which never experienced starvation stress, contained a relatively high percentage of cells carrying low-level amplifications of the *cat* genes, likely duplications (two copies). Again, this is evident in figure 7C by either enlarged NotI E fragments (Pre. 3) or an additional NotI fragment of relatively low intensity (Pre. 1 and 2). In each of these three precursor populations, these PFGE results show a significantly higher percentage of cells with low-level *cat* gene amplifications than unenriched Ben-parental DR1023 populations which typically do not have PFGE-detectable amplifications or other rearrangements.

Notably, each precursor had approximately the same amplicon size as its three revertant siblings. The Rev. 3ABC sibling set with enlarged NotI E fragments had a precursor (Pre. 3) with an E fragment enlarged by approximately 40 kb, consistent with a duplication. Corroborating evidence supporting Precursor 3 carries a duplication is provided by its three sibling revertants having a ladder of various size enlarged E fragments where each rung is separated by approximately 40 kb, suggesting the amplicon size is ∼40 kb. Similarly, the Rev. 1ABC and Rev. 2ABC sibling sets with an additional NotI fragment (equivalent to the amplicon size) had precursor populations with the same amplicon size: 237 kb in Pre. 1 and 247 kb in Pre. 2 (fig 7). In these two cases, the additional NotI fragment had a much lower relative intensity in the precursor populations, indicating a lower amplicon copy number compared to their revertant sibling populations. This lower band intensity also likely reflects the higher instability and collapse of their larger duplicated segments compared to Pre. 3 during the nonselective growth. Together, these results indicate Ben+ high-copy amplification mutants arise from gene duplication events that occur prior to selective stress.

### Independent populations show significant fluctuation in amplification mutant frequencies rather than a Poisson distribution

Our experimental results from two independent methods, replica plating and reconstruction assays, suggest Ben+ revertants originate from rare duplication events that occur during the LB growth before selective pressure. As a third independent method to either support or oppose these findings, we performed a Luria-Delbrück fluctuation test to determine whether these mutants arise before or during selection. The basis of this test is that mutant frequencies vary or “fluctuate” between different parental cultures due to random early or late mutational events occurring during cell growth before selection. In the Ben reversion, if ancestral precursor mutants arise during nonselective growth prior to starvation stress, they should have Ben+ mutant frequencies that widely fluctuate between independent LB cultures. In contrast, the adaptive amplification theory argues mutant frequencies should not fluctuate between cultures since stress induction of mutant formation is proposed to occur on the selective plates, where nutrient limitation and growth restriction occurs. If mutations are induced at fixed rates during selection, where parental population sizes are relatively constant and do not change much, there should be very little to no fluctuation in Ben+ mutant frequencies between different cultures (other than fluctuation caused by procedural or sampling variation).

For our fluctuation test, we grew 119 parallel independent LB cultures of the Ben-parental DR1023 strain, with each culture inoculated from a different single isolated colony. After overnight growth, one sample from each culture was spread onto a single selective Ben plate. As a control to analyze intraculture fluctuation (due to procedural or sampling variation), a large DR1023 culture was spread onto many selective plates. To access the repeatability of this control, 3 large DR1023 cultures were spread onto a total of 116 selective plates (18, 49, and 49). All 235 selective plates were incubated at 30°C and analyzed every day for 21 consecutive days. Our fluctuation test results show Ben+ revertant colonies accumulated on the selective plates at the same average rates both in the single and multiple cultures, as shown in figure 3. However, “jackpots” were seen amongst only the 119 independent cultures, and not seen in any of the 116 selective plates from the 3 large cultures (table 1). For example, 2 of the 119 selective plates from independent cultures had 17 and 19 revertant colonies per Ben plate on day 10, when the median was 1 Ben+ revertant colony per plate. Another 4/119 plates had 10, 10, 10, and 12 Ben+ revertant colonies per plate. In comparison, none of the 116 plates derived from the 3 large cultures had over 2 Ben+ revertant colonies per plate on day 10.

**Table 1.** Luria and Delbrück fluctuation test results showing the distributions of Ben+ gene amplification mutant frequencies between and within cultures. For the 119 cultures, one sample of each culture was plated onto a single Ben plate. For the 3 large cultures, each culture was plated onto multiple Ben plates: 18, 49, and 49. Mutant frequencies were measured by the number of Ben+ revertant colonies per Ben plate after 10 days of incubation. The mutant frequency distributions in the 119 parallel cultures are shown in the column to the left of the vertical line, while each of the 3 large cultures are shown in the 3 columns to the right.

|  |  |  |  |  |
| --- | --- | --- | --- | --- |
| **Number of cultures** | 119 | 1 | 1 | 1 |
| **Number of plates** | 119 | 18 | 49 | 49 |
| **Number of Ben+ revertants per plate** |  | **Number of plates** |  |  |
| 0 | 48 | 10 | 40 | 39 |
| 1 | 24 | 4 | 6 | 9 |
| 2 | 21 | 4 | 3 | 1 |
| 3 | 7 |  |  |  |
| 4 | 8 |  |  |  |
| 5 | 1 |  |  |  |
| 6 | 1 |  |  |  |
| 7 | 1 |  |  |  |
| 8 | 1 |  |  |  |
| 9 | 1 |  |  |  |
| 10 | 3 |  |  |  |
| 11 |  |  |  |  |
| 12 | 1 |  |  |  |
| 13 |  |  |  |  |
| 14 |  |  |  |  |
| 15 |  |  |  |  |
| 16 |  |  |  |  |
| 17 | 1 |  |  |  |
| 18 |  |  |  |  |
| 19 | 1 |  |  |  |
| **Average per plate** | 1.95 | 0.67 | 0.24 | 0.22 |
| **Variance** | 10.13 | 0.71 | 0.31 | 0.22 |
| **Median** | 1 | 0 | 0 | 0 |

Notably, for all days analyzed there was much higher variation in Ben+ mutant frequencies per Ben plate between multiple cultures than within single cultures, for all three large cultures (fig 8A). The variance was 10- to 190-fold higher in multiple cultures compared to single cultures depending on the day and culture. A Brown-Forsythe equality of variance test indicated a significant difference between the variance in multiple versus single cultures. This was true for all days analyzed except for four days in one of the three large cultures; yet these four days still had over 19-fold higher variance across the multiple cultures than within the single culture. Together, these results indicate the high fluctuation observed in the 119 parallel cultures was not due to procedural or sampling variation. Importantly, the high variance between multiple cultures for all days further suggests that Ben+ gene amplification mutants originate before selection, including revertants arising late.

**Figure 8.**
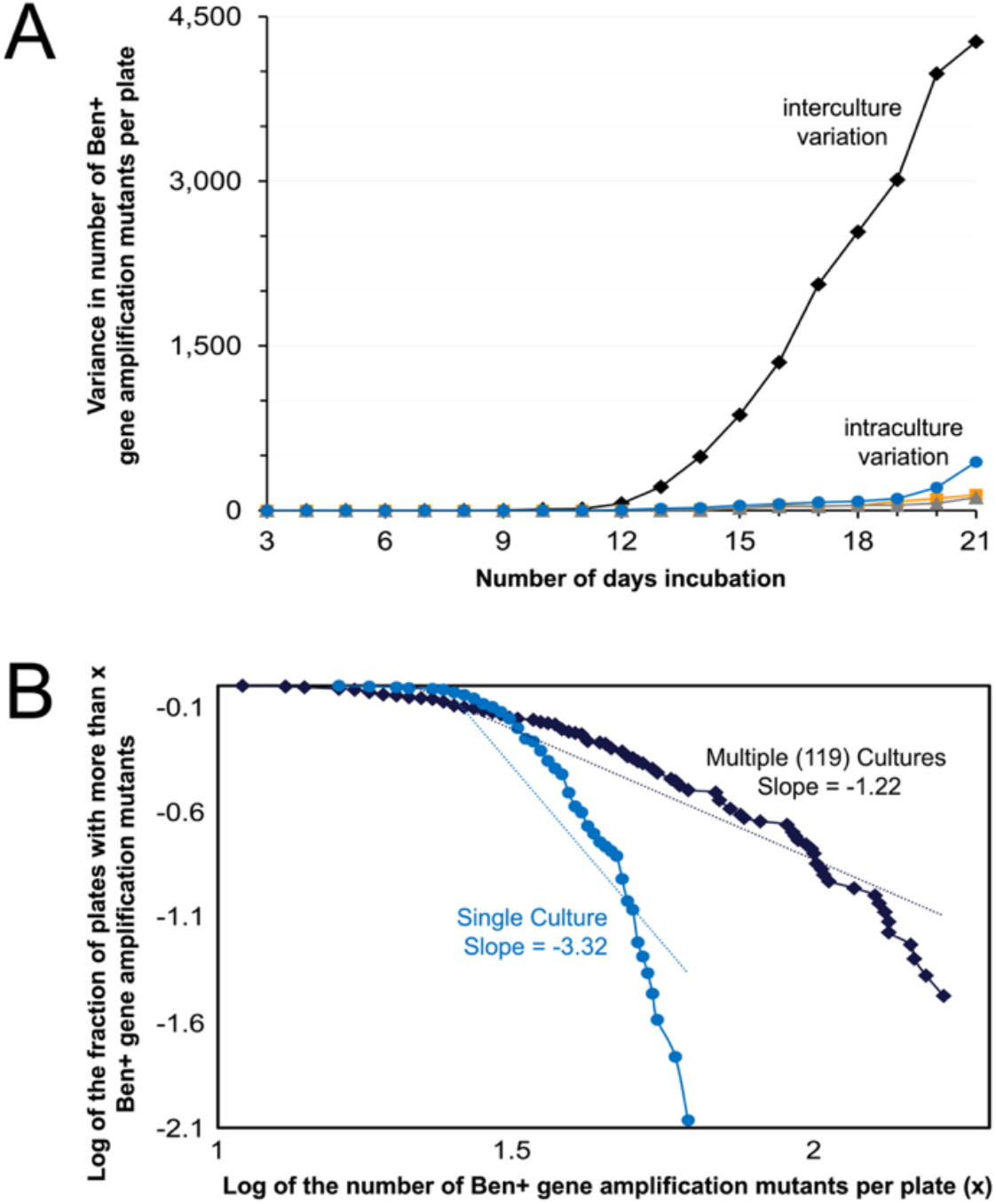
Fluctuation test results comparing Ben+ mutant frequencies in multiple cultures to single cultures. A) The amount of variance in the Ben+ mutant frequencies per Ben plate in parallel cultures compared to single cultures, for all days analyzed. The black diamonds refer to the amount of variance between 119 parallel cultures where one sample of each culture was spread onto a single Ben plate. The orange squares, gray triangles, and blue circles each refer to the amount of variance within a single culture where each culture was spread onto 18, 49, and 49 Ben plates, respectively. B) The distributions of Ben+ revertants per Ben plate in multiple and single cultures after 19 days of incubation. The y-axis indicates the observed number of plates with more than x Ben+ colonies for the various values of x shown on the x-axis. The black diamonds show to the distribution across the 119 parallel cultures described in panel A, whereas the blue circles show to the distribution within a single culture spread onto 49 selective plates. The slopes are based on the best fit dotted line shown in blue and black.

Furthermore, we analyzed the distribution of Ben+ mutant frequencies per Ben plate across the independent cultures to determine whether they follow a Poisson or Luria-Delbrück fluctuation distribution. The adaptive amplification theory predicts a Poisson distribution (little to no fluctuation) in both the single and multiple cultures. In contrast, the view that gene amplification mutants originate before selection predicts a Poisson distribution in single cultures, but a Luria-Delbrück fluctuation distribution (wide fluctuation) across multiple cultures. Mutant distributions were analyzed by plotting the relationship between the logarithm of the proportion of selective plates with *x* or more Ben+ mutants (y-axis) and log *x* (x-axis) (Luria and Delbrück 1943; Cairns et al. 1988; Cairns and Foster 1991). As shown in figure 8B, our analysis demonstrates the three single cultures each followed a Poisson distribution across the multiple Ben plates, as evident by a relatively flat line followed by a steep drop (slope = −3.32). This Poisson distribution indicates very little fluctuation in Ben+ mutant frequencies measured on different selective plates derived from a single culture. In comparison, the multiple cultures followed a mix of Luria-Delbrück fluctuation and Poisson distributions. Compared to the distribution within single cultures, the distribution across multiple cultures is much closer to a straight line (slope = −1.22), indicating a Luria-Delbrück fluctuation distribution (fig 8B). This distribution suggests the Ben+ gene amplification mutants originated prior to selection. However, this line has a moderate downward inflection, consistent with a partial Poisson distribution. Although these mixed distributions are difficult to directly interpret, they are consistent with our results demonstrating Ben+ revertants are derived from duplication mutations formed before selection. Unlike point mutants, duplication mutants rapidly approach a steady-state frequency during nonselective growth due to their high loss rate (collapse to haploid) and associated fitness cost (Reams et al. 2010). Even if the first precursor duplication forms early or late during the LB growth prior to selection, all independent cultures approach the same *cat* gene duplication steady-state frequency, thereby limiting fluctuation (Reams and Roth 2015). This could explain the observed partial Poisson distribution. Still, the distribution across independent cultures was closer to a Luria-Delbrück fluctuation distribution than the Poisson distribution in single cultures. In summary, our fluctuation test results showing multiple cultures have jackpots, significant high variance, and a trend towards a Luria-Delbrück fluctuation distribution further supports that Ben+ gene amplification mutants originate before selective stress.

## Discussion

DNA-damaging agents — such as radiation and DNA reactive chemicals — increase mutation rates through error-prone DNA damage repair responses (e.g., SOS, etc.) (Maslowska et al. 2019). However, it is still debated whether growth inhibition caused by selective environments triggers mutagenesis without DNA-damaging agents present. Here, we tested the adaptive amplification theory, proposing amplification mutations are induced by selective stress, using an *Acinetobacter* system that exclusively analyzes amplification mutants. Our experimental results demonstrate amplification mutants originate from low-copy amplifications (duplications) that arise before exposure to selection; therefore, the initial duplication cannot be caused by selective stress. Indeed, gene duplication mutations form frequently without selection in diverse organisms and contribute to many known human neurological disorders (Lupski et al. 1991; Flores et al. 2000; Zhang et al. 2009; Ramocki et al. 2010; Reams et al. 2012; Reams et al. 2014; Konrad et al. 2018). In an unselected lab culture of *Salmonella enterica,* approximately 10% of cells carry a duplication of some chromosomal region (Roth et al. 1996). In an *A. baylyi* overnight nonselective culture, Seaton and coworkers measured *cat* gene duplications at a frequency of 0.01%, equivalent to about 50,000 cells per benzoate selective plate in the Ben reversion assay (Seaton et al. 2012). This number of plated *cat* duplication bearing cells (50,000) is vastly greater than the number of revertant colonies (70) after 3-weeks incubation, suggesting that the plated *cat* duplications have a low probability of reaching visible colonies. However, it should be noted that Ben+ revertant colonies continue to accumulate on the selective plates beyond 3 weeks. Also, while different Ben+ revertants carry *cat* gene amplicons that vary widely in size (12-290 kb), one of the amplicon ends appears to be restricted to an approximate 5.5 kb region between the *catA* gene and the upstream promoter of the *benABCDE* operon (Reams and Neidle 2004a). This endpoint limitation may reduce the number of *cat* duplications that can amplify to support benzoate growth. Even with these complexities, we estimate there is a sufficient number of cells carrying permissive *cat* gene duplications present before selection to account for the number of revertants.

Although the initial duplication appears to arise before selection, the secondary steps of generating higher-copy *cat* gene amplification occur during selection. Still, these secondary steps do not appear induced by selective stress for several reasons. First, our reconstruction experiments with different copy number strains indicate these higher amplification steps occur relatively slowly under selection, over many days to multiple weeks; therefore, they do not appear to be induced. Second, our fluctuation test results show the number of new revertants arising on all days, even 21 days, highly fluctuate between independent cultures compared to single cultures, suggesting they originated before selection.

Rather than stress-induced, Ben+ high-copy amplification colonies can be explained by a clonal evolution process driven by natural selection, consistent with the amplification under selection model proposed by Roth (fig 9). According to this model, rare cells with a *cat* duplication are present before selection. On the selective plate, these cells initiate slowly growing clones. Within the developing clones, selection favors cells with further increases in *cat* copy number (higher amplification). These likely form by unequal recombination between the amplified segments (repeats), through the same mechanisms as their collapse (fig 1). Since the substrates of these events are large repeats (12-290 kb), these events likely occur spontaneously at sufficiently high rates (∼10^-2^ / cell / generation), providing a mathematical practicality for generating amplification within the time it takes to form a visible colony (∼20 generations) (Reams et al. 2010). Before selection, higher amplification is limited due to frequent collapse and associated fitness costs. However, under selection, copy number increase is driven since selection favors growth of cells with more *cat* gene copies (fig 9).

**Figure 9.**
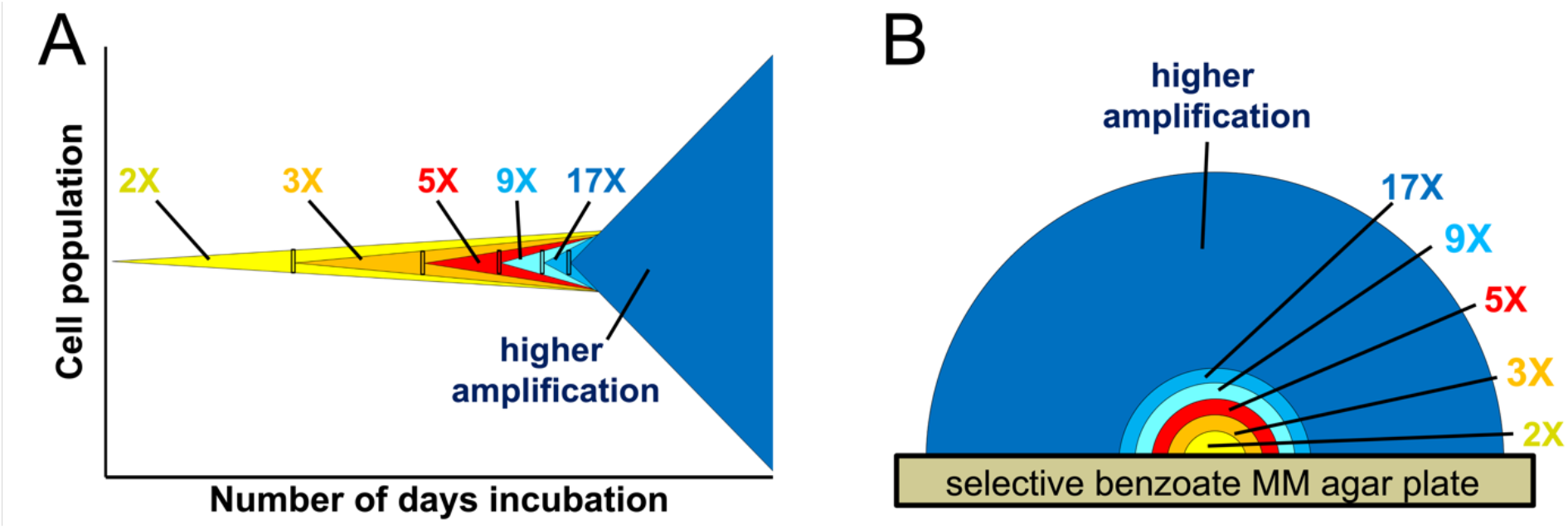
Clonal expansion of precursor mutants during selection. A) The x-axis shows the number of days of incubation (increasing from left to right) during selection and the y-axis represents the parental cell population. On day 0 (far left x-axis), a nonselectively grown parental cell population is plated onto growth restrictive selective Ben media. Within this population, rare variants carrying tandem duplication mutations encompassing the *cat* genes (2X) are present at a frequency of approximately 0.01% of the cell population. During incubation, selection provides a growth advantage to these variants (due to their increased ability to degrade benzoate) allowing them to slowly outcompete the haploid parental cells over time (thereby forming microscopic colonies on selective plates). Once the *cat* duplication-carrying cells accumulate to approximately 100 cells (indicated by the small yellow-filled rectangle), they are likely to contain higher amplification variants with three copies (3X). Selection favors these higher copy variants allowing for a series of higher amplification events within a single clonal population, eventually forming a visible high amplification revertant colony. B) Formation of a Ben+ revertant colony. The amplification model by Roth and coworkers predicts each colony should include representatives of the initial precursor and each higher amplification step, since individual improved cells grow faster but retain their unimproved ancestors within the population (Roth et al. 2006).

This process of clonal evolution, where a “renegade cell” expands and diversifies through sequential mutational events, plays an important role in cancer development, therapy resistance, and relapse (Andor et al. 2016; McGranahan and Swanton 2017; Weinberg 2008). Also, amplifications are found in many tumors where amplified genes (e.g., *MYC*, *CCNE1, HER2*) help cells escape growth limitation (Ahn et al. 2020; Gorski et al. 2020). Based on our results, we suggest duplications of growth factor genes may serve as genomic biomarkers for early cancer detection and treatment, before high-copy amplification is attained (Caporaso 2013). For example, in several reported cases of metastatic breast cancer, *MYC* duplications in primary tumors were found to expand to higher-copy *MYC* amplification in the metastatic more aggressive breast cancers (Singhi et al. 2012). In the future, understanding and preventing the mechanisms of generating higher copy number will be important for targeting and inhibiting amplification in cancer cells.

## Data availability

Strains are available upon request. The authors affirm that all data necessary for confirming the conclusions of the article are present within the article, figures, and tables.

## Acknowledgements

We are grateful for the support of our staff, especially Douglas Whited, Richard Aguirre, Nancy Burford, and Gordon Zanotti. We thank John Roth for helpful suggestions.

